# Lateralized deficits in arousal processing after insula lesions: behavioral and autonomic evidence

**DOI:** 10.1101/2021.03.24.436828

**Authors:** Olga Holtmann, Marcel Franz, Constanze Moenig, Jan-Gerd Tenberge, Insa Schloßmacher, Iskrena Ivanova, Christoph Preul, Wolfram Schwindt, Nico Melzer, Wolfgang H.R. Miltner, Thomas Straube

## Abstract

A large body of evidence ascribes a pivotal role in emotion processing to the insular cortex. However, the complex structure and lateralization of emotional deficits following insular damage are not understood. Here, we investigated emotional ratings of valence and arousal and skin conductance responses (SCR) to a graded series of emotionally arousing scenes in patients with left (*n* = 10) or right (*n* = 9) insular damage and in healthy controls (*n* = 18). We found a significant reduction in overall SCRs, arousal ratings and valence extremity scores in right-lesioned patients, as compared to left-lesioned patients and healthy controls. Additional analyses of correlations between subjective arousal ratings resp. SCR and normative arousal ratings revealed that both lesion groups had evaluative and physiological difficulties to discover changes in stimulus arousal. Although no group differences emerged on overall ratings of valence, analysis of correlations between subjective and normative valence ratings displayed markedly reduced accuracy in right-lesioned patients, as compared to left-lesioned patients and healthy controls. Our findings support the hypothesis that the left and right insulae subserve different functions in emotion processing, potentially due to asymmetrical representations of autonomic information in the left and right human forebrain. The right insula may serve as integral node for sympathetic arousal and cognitive affective processing.

## 1. Introduction

The insular cortex plays a central role in emotion processing (Craig, 2014; Gu et al., 2013; Menon & Uddin, 2010). A substantial body of functional neuroimaging and electrophysiological research has demonstrated insular involvement in interoceptive awareness and autonomic regulation, subjective feeling, and motor expression of emotion (Adolfi et al., 2017; Dondaine & Péron, 2012; Lamm & Singer, 2010; Lindquist et al., 2012, 2016; Mai et al., 2019; Zaki et al., 2012). Insula damage of various etiologies has been associated with impaired evaluation of emotional valence and arousal (Berntson et al., 2011), disrupted experience and recognition of emotion in self and others (Kumfor et al., 2017), in apathy, emotional blunting, and autonomic dysregulation (for reviews see Ibañez et al., 2010; Jones et al., 2010; Vicario et al., 2017). However, it is unclear whether some emotional processes rely more on the insula than others. Some lines of research suggest a major involvement in decoding emotional valence (especially for negative emotions) (Gogolla, 2017; Kayyal et al., 2019), while recent models propose a disproportionate role in interoception and emotional arousal (Craig, 2014; Critchley, 2009; Jones et al., 2010). Thus, it remains to be investigated in detail, how insula damage refers to psychological frameworks, which conceptualize emotion along two dimensions, valence (level of experienced pleasantness/ unpleasantness) and arousal (level of intensity of a given emotion), that are mirrored in differential physiological responses (Bradley & Lang, 2007; Lang, 2014). In particular, skin conductance response (SCR) is considered a reliable autonomic marker of emotional arousal and its somatovisceral impact (Lang, 2014).

Furthermore, emotional functions are increasingly recognized being lateralized in the human brain, though the exact nature of affective asymmetries is highly debated (Gainotti, 2018). Given the broad role of the insula in emotion, the question of laterality effects within the insula is intriguing. Functional imaging and electrophysiological research have suggested a superior role of the right insula in autonomic and emotional functions (Craig, 2002; Critchley, 2009). Accordingly, some clinical studies have shown associations between right insula lesions and disturbed evaluation of emotion by means of voxel-based morphometry techniques (Adolphs et al., 2000; Tippett et al., 2018). A few studies compared patients with asymmetrical forms of fronto-temporal degeneration including the insula and found more severe emotional abnormalities in patients with right atrophic predominance (Binney et al., 2016; Irish et al., 2013). Terasawa et al. (2015) reported attenuated sensitivity for negative emotions in three cases of right insular, frontal and temporal damage. Couto et al. (2013) documented deficits of multimodal aversive emotion recognition, empathy and contextual inference of emotions in a patient with focal right-hemispheric damage of white matter association tracts adjacent to the insula. Kim et al.(2017) showed emotional blunting, as assessed by ratings and SCR, to wins and losses in a roulette game in a group of nine patients with right insular lesions. However, functional neuroimaging studies also report left (Small et al., 2003; Wicker et al., 2003), or bilateral (Vytal & Hamann, 2010) insula involvement in emotion. A study of patients with traumatic brain injury showed that the left insula, as part of a bilateral fronto-temporo-limbic network, is involved in emotion recognition from facial expressions (Dal Monte et al., 2013). Left insula atrophy has been linked to impaired disgust recognition in pre-symptomatic Huntington’s disease patients (Henley et al., 2008; Kipps et al., 2007), and frontotemporal disease patients (Kumfor et al., 2013). Single case studies of patients with left insula damage unveiled selective impairments of recognition and experience of disgust (Borg et al., 2013; Calder et al., 2000). In a previous study, we found a selective disgust deficit in a group of four patients with left insula damage, as compared to a group of four patients with right insula damage, on emotion-specific composite scores (Holtmann et al., 2020). It has been suggested that emotional processing is left- or right-lateralized based on differential autonomic inputs to the human forebrain, with parasympathetic representations in the left, and sympathetic representation in the right hemisphere (Craig, 2014). However, there is a lack of studies that systematically investigated arousal and valence responses in patients with lateralized insula lesions. Thus, the roles of the right and left insulae in emotion processing are not clear. Systematic comparisons of patients with right- and left-lateralized brain damage matched for size and distribution are needed to clarify potentially distinct hemispheric contributions to valence and arousal processing.

In the current study, we investigated the effect of lateralized insula damage on valence and arousal ratings and autonomic responses to visual emotional stimulation. We examined ten patients with left and nine patients with right insula damage who were matched on demographic and clinical parameters. Eighteen subjects matched for age and educational background with no history of psychiatric or neurological conditions served as healthy controls. Given the role of the insula (and especially the right insula) in interoception and arousal functions (Critchley, 2005), we assumed we would find disproportionate changes in measures of arousal, particularly after right-hemispheric lesions.

## 2. Materials and methods

### 2.1. Participants

Nineteen patients (nine females) with a history of either left (*n* = 10) or right (*n* = 9) damage centrally affecting the insular cortex were enrolled in the study. Patients were recruited to the research programme through the Department of Neurology, University Hospital Muenster, Muenster, Germany (*n* = 7), or following participation in a rehabilitative programme (Miltner et al., 1999, 2016) offered by the Department of Clinical Psychology, Friedrich Schiller University Jena, Jena, Germany and the Department of Neurology, University Hospital Jena, Jena, Germany (*n* = 12). Inclusion criteria were as follows: (1) unilateral lesions due to middle-cerebral-artery stroke, centrally affecting the insular cortex, as confirmed by three experts in clinical neuroimaging (CP, WS and NM) from CT or MRI scans of the brain; (2) stable lesions (at least 1 year after lesion onset); (3) no cognitive deficits compromising the understanding of task instructions and task performance (i.e., global aphasia, attention deficits, amnesia, disorders of reasoning, or visual neglect); (4) no history of neurodegenerative disorders, epilepsy, brain tumours, or brain trauma; (5) no history of substance-induced disorders; (6) no history of psychiatric disorders. Clinical and demographical data are reported in Table 1.

**Table 1.** Sample characteristics.

|  | Left-lesioned<br>group<br>*n* = 10 | Right-lesioned<br>group<br>*n* = 9 | Control<br>group<br>*n* = 18 | Statistical<br>comparison<br>(*P*) |
| --- | --- | --- | --- | --- |
| **Demographic** |  |  |  |  |
| Age (years) | 58.0 (1.7) | 59.4 (2.7) | 61.6 (1.6) | 0.382 |
| Gender (males/ females) | 6/4 | 4/5 | 10/8 | 0.781 |
| Education (years) | 12.6 (0.3) | 11.2 (0.7) | 11.7 (0.4) | 0.213 |
| **Clinical** |  |  |  |  |
| Lesion age (years) | 6.5 (1.0) | 5.1 (1.0) | N/A | 0.342 |
| Lesion volume (cm<sup>3</sup>) | 41.2 (16.9) | 45.7 (19.1) | N/A | 0.860 |
| **Neuropsychological (z-scores)** |  |  |  |  |
| IQ (WAIS-IV-MR) | 0.4 (0.3) | 0.2 (0.3) | N/A | 0.748 |
| Visuo-spatial Memory (CFT) | 0.5 (0.3) | 0.4 (0.3) | N/A | 0.777 |
| Auditory Memory<br>(immediate recall; WMS-IV-LMi) | -0.6 (0.3) | -0.7 (0.3) | N/A | 0.938 |
| Auditory Memory<br>(delayed recall; WMS-IV-LMd) | -0.4 (0.2) | -0.7 (0.4) | N/A | 0.640 |
| Phasic Alertness (TAP-Alert) | -0.3 (0.4) | -0.8 (0.3) | N/A | 0.289 |
| Selective Attention (TAP-Go/NoGo) | -0.04 (0.2) | -0.1 (0.2) | N/A | 0.980 |
| Divided Attention (TAP-DA) | -0.1 (0.3) | -0.02 (0.5) | N/A | 0.929 |
| Visual Field/ Neglect<br>(TAP-VFCT, blind spots) | 0 | 0 | N/A | -- |
| **Affective Screening** |  |  |  |  |
| BDI-II | 14.6 (2.5) | 11.0 (2.6) | 3.3 (0.6) | < 0.001 |

*Note:* Data are means (standard error of mean), except for gender (absolute frequencies). Statistical comparisons: one-way ANOVA, chi-squared test (gender). BDI-II = Beck Depression Inventory, CFT = Rey Complex Figure Test, TAP = Test Battery for Attention Performance, TAP-Alert = TAP-subtest 'Alertness', TAP-Go/NoGo = TAP-subtest 'Go/NoGo', TAP-DA = TAP-subtest 'Divided attention I, aud-vis', TAP-VFCT = TAP-subtest 'Visual field with central task', WAIS-IV-MR = subtest 'Matrix Reasoning' from the Wechsler Adult Intelligence Scale (4<sup>th</sup> edition), WMS-IV-LMi = subtest 'Logical Memory', immediate recall, from the Wechsler Memory Scale (4<sup>th</sup> edition), WMS-IV-LMd = subtest 'Logical Memory', delayed recall, from the Wechsler Memory Scale (4<sup>th</sup> edition)

Eighteen neurologically and psychiatrically healthy controls (eight females) of similar age and educational background were recruited via public ads in Muenster (*n* = 8) and Jena (*n* = 10). All subjects gave informed consent for participation in the research in accord with the Declaration of Helsinki and the positive decision for the present study by the Ethics Committee of the German Society of Psychology (TS032013_rev). All participants were right-handed.

### 2.2. Lesion reconstruction

High-resolution isotropic (1mm) 3D structural T_1_-weighted anatomical images were acquired on a 3 Tesla system (Magnetom Prisma, Siemens, Erlangen, Germany) for 15 patients within 40 days after participation. Four patients did not provide consent for MRI or did not meet the inclusion criteria for MRI at the time of testing. In these cases, lesion analyses were based on T_*2*_FLAIR-weighted MR images (3mm) obtained at 1.5 Tesla (Philips Achieva, Philips Medical Systems, Best, The Netherlands; *n* = 3) or cranial CT images (5 mm; Somatom Definition Flash Siemens Medical Solutions, Forchheim, Germany; H30s soft kernel; *n* = 1) from the latest clinical routine follow-up screening. The anatomical images were aligned to Montreal Neurological Institute (MNI) space (1 x 1 x 1 mm^3^) by means of the Clinical Toolbox for SPM12 (Wellcome Trust, London, UK), which includes standard stroke-control MRI and CT templates based on healthy older participants and normalization procedures recommended for stroke-aged populations (Rorden & Brett, 2000; Rorden et al., 2012). Individual lesion boundaries were identified by means of the Clusterize Toolbox (version 1.0beta) available for SPM12, offering a semi automatic approach to lesion demarcation regardless of image modality (Clas et al., 2012; de Haan et al., 2015). During the automated pre-processing procedure, each image was inspected for local intensity maxima, thus defining cluster cores. Minimum size for the initiation of a cluster was set to 150 mm^3^ and a lower intensity threshold of 20% was used for the elimination of irrelevant background voxels (de Haan et al., 2015). The intensity threshold was then iteratively adapted in steps of 1%, resulting in the assignment of each voxel to a cluster core. Clusters corresponding to damaged tissue were then interactively selected and, if necessary, modified by changing the intensity threshold. Finally, resulting lesion demarcations were reviewed and manually adjusted by a professional neurologist (CM) who was blind to the study’s hypotheses, data and statistical analyses at this time. MriCron (http://www.mccauslandcenter.sc.edu/crnl/tools) was used to create within-group overlaps and display them on a T_1_-weighted MNI template (Rorden et al., 2007). Regional extent of damaged tissue was determined by using standard anatomical atlases implemented in MriCron, i.e. Automated Anatomic Labelling (AAL) atlas (Tzourio-Mazoyer et al., 2002) and JHU white matter tractography atlas (Oishi et al., 2008).

### 2.3. Experimental design

To study valence and arousal independently, we created a stimulus set following the approach by Glaescher and Adolphs (2003). Forty-four emotionally charged color images were drawn from the International Affective Picture System (Lang et al., 2008). The stimuli were selected to broadly cover the valence-arousal affective space according to their valence and arousal norms (see Fig. 1). Pleasant and unpleasant stimuli of high, medium, and low arousal, covering a broad range of emotional content and social as well as non-social contexts, were included. Neutral stimuli of low and medium arousal complemented the distribution. We explicitly excluded disgust stimuli to avoid biases in emotional processing since there is evidence that the insula may differentiate disgust from all other basic emotions (Calder et al., 2000; Holtmann et al., 2020; Vicario et al., 2017). Using Adobe Photoshop, mean luminance was aligned between categories of different emotional content. All images were presented in landscape.

**Figure 1.**
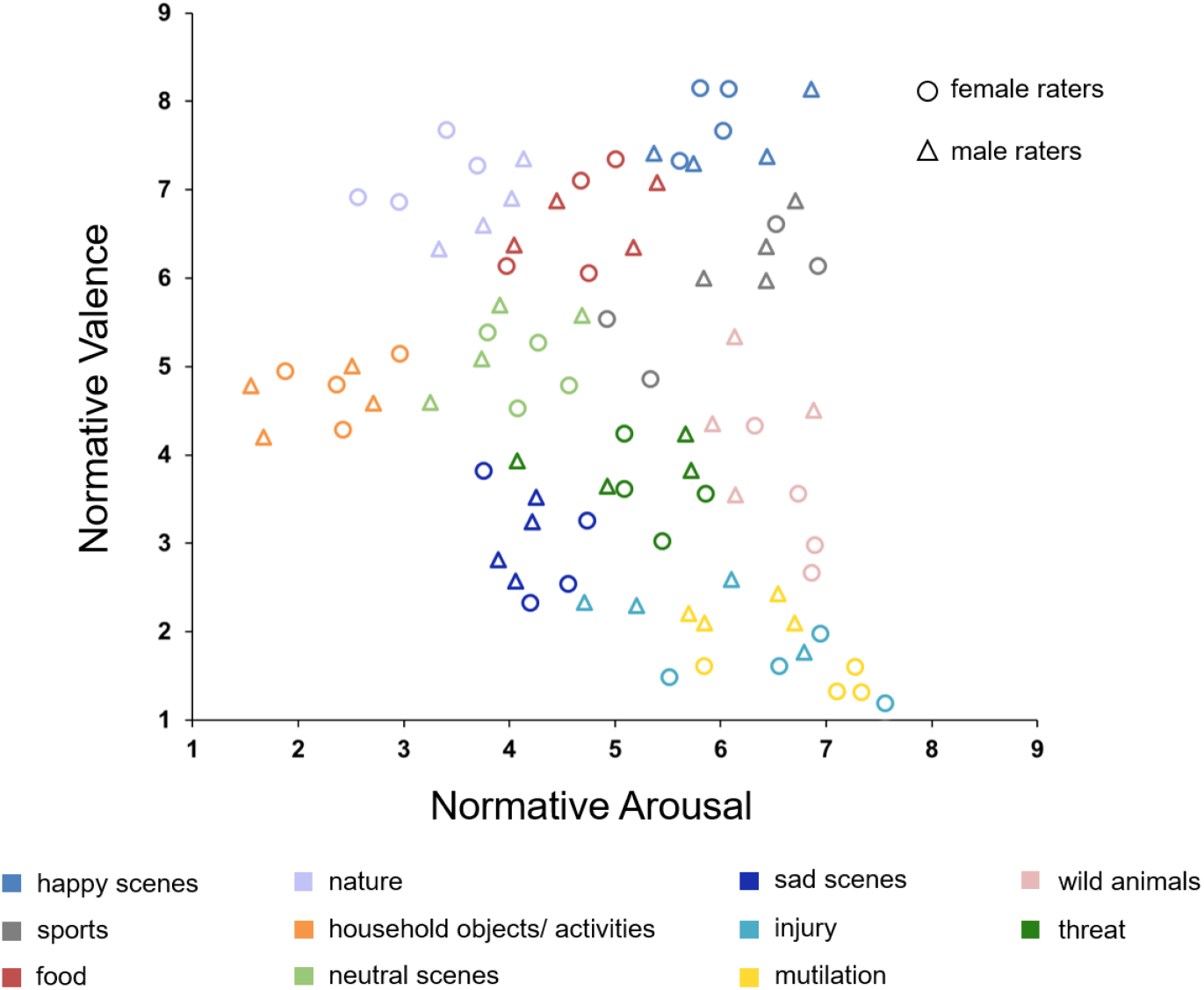
Dimensional representation of the 44 selected images based on normative arousal and valence ratings, as provided by Lang and colleagues (Lang et al., 2008). Content categories are highlighted in different colors. Note that higher valence scores correspond with more pleasant emotional experience. Higher arousal scores indicate more arousing emotional experience.

The study protocol was identical for measurements in Muenster and Jena. Each of the 44 images was presented for 5000 ms in the center of a 27 inch computer screen on a black background in a dimly lit room. Seating distance from the screen was approx. 60 cm. Images were presented in randomized order, with no repetition of images belonging to the same emotion content category. Subjects received standardized instructions to watch the scenes carefully, and by using nine-point Likert scales to indicate subjective valence (−4 = *very negative*, 0 = *neutral*, 4 = *very positive*) and arousal (1 = *not at all arousing*, 9 = *highly arousing*) after each scene. Both rating tasks were self-paced, but subjects were encouraged to respond as quickly as possible. The rating tasks were followed by a fixation cross with a jittered duration of 5600 – 6400 ms, *M* = 6000 ms, allowing the electro-dermal activity to return to baseline (Boucsein et al., 2012; Dawson et al., 2007; Figner & Murphy, 2011). Participants were trained on the rating tasks using two practice images with neutral content that were not included in the stimulus set. Stimulus presentation and response recording were controlled by Presentation Software (v19.0, Neurobehavioral Systems, Albany, California, USA). Task performance was addressed by means of valence, arousal, and extremity scores. Extremity scores were computed as the absolute value of the valence rating minus the midpoint of the scale (i.e. *abs*[valence -0]) and were added to improve the grasping of potential response biases in the patient groups.

### 2.4. Neuropsychological functions

All patients underwent a detailed neuropsychological evaluation targeting major neurocognitive domains prior to participation. General intelligence was measured by the subtest ‘Similarities’ from the German adaptation of the Wechsler Adult Intelligence Scale (Daseking & Petermann, 2013). Visuo-spatial memory was assessed by the Rey Complex Figure Test (Meyers & Meyers, 1995), auditory memory (immediate and delayed recall) was assessed by the subtest ‘Logical Memory’ from the Wechsler Memory Scale (Lepach & Petermann, 2012). Visuospatial attention was addressed by means of the Test Battery for Attention Performance (Zimmermann & Fimm, 1997) (subtests ‘Alertness’, ‘Go/NoGo’, ‘Divided Attention I, aud-vis’ and ‘Visual field with central task’, respectively addressing phasic alertness, selective attention, divided attention, and visual field defects due to neglect). Severity of depression symptoms at the time of testing was assessed by the Beck Depression Inventory (BDI-II) (Beck et al., 1996) in patients and controls.

### 2.5. Skin conductance recording and pre-processing

During the entire duration of the protocol, skin conductance was acquired by self-adhesive disposable Ag/Ag-Cl surface electrodes (8 mm diameter, filled with conductive electrode gel) placed on the thenar and hypothenar eminence of both hands. The skin conductance signal was transmitted to an ECG100C amplifier (BIOPAC Systems, Inc., Goleta, California, USA) in Muenster, and to an UBV 400 SC coupler and amplifier (Rimkus Medizintechnik, München, Germany) in Jena. Both amplifiers used a constant voltage (0.5 V) technique to measure skin conductance. Skin conductance signals were received by the BIOPAC AcqKnowledge software (v4.3) in Muenster, and by the DASYLab software (v5.0, Data Acquisition System Laboratory, Mönchengladbach, Germany) in Jena. Raw signals were digitized at 1000 Hz in Muenster, and at 500 Hz in Jena. The room temperature in both labs was kept similar (approx.: 21°C). Before starting the experimental investigation of emotional processing, a standardized respiration measurement was conducted (Holtmann et al., 2018; Schröder et al., 2015), during which the general ability to develop SCRs was proved. Only after passing this pre-test, the experiment started. Raw data were transformed using the BIOPAC AcqKnowledge software in Muenster, and the Brain Vision Analyzer software (v2.1, Brain Products GmbH, München, Germany) in Jena for analysis in Matlab (v9.1, The MathWorks, Inc., Natick, Massachusetts, USA). After careful visual inspection, data from one control subject and two right-lesioned patients were discarded from analysis due to severe artefacts. The final dataset included 34 subjects (*n* = 17 controls [*n*_*Muenster*_ = 7, *n*_*Jena*_ = 10, seven females], *n* = 10 left-injured patients [*n*_*Muenster*_ = 3, *n*_*Jena*_ = 7, four females], and *n* = 7 right-injured patients [*n*_*Muenster*_ = 4, *n*_*Jena*_ = 3, three females]). The data were down-sampled to 250 Hz. A high-pass filter of 0.5 Hz was applied to remove tonic changes; a low-pass filter of 2 Hz was used to remove high-frequency noise. The area under the curve (AUC), a common indicator of the SCR (Boucsein et al., 2012; Figner & Murphy, 2011; Gläscher & Adolphs, 2003), was extracted as the positive time integral of the response amplitude from the band-pass filtered signal in a 5 sec interval starting with the presentation of each stimulus (Gläscher & Adolphs, 2003). The AUC scores were log-transformed for usage in statistical analyses (Boucsein et al., 2012). We observed no significant differences between recordings from both hands (controls: *P* = 0.348, left-lesioned group: *P* = 0.901, right-lesioned group: *P* = 0.367). Therefore, we report pooled results from both hands.

### 2.6. Statistical analyses

Lesion groups were contrasted on clinical and neuropsychological variables by means of two-sided two-sample *t*-tests. Group differences between lesion groups and the control group in demographic variables and mood were assessed using one-way ANOVAs for continuous variables and chi-square analyses for categorical variables.

To assess the effects of lateralized insular damage on the processing of emotional stimuli, we pursued a two-step approach (see Gläscher & Adolphs, 2003). First, we compared the groups (left-lesion group, right-lesion group, healthy control group) on mean valence, extremity, arousal and AUC scores, thus exploring potential group differences in terms of response levels. Second, we focused on the normative ratings as a systematic scale of comparison to address group differences. We correlated subjective rating data and AUC data with normative valence and arousal ratings (Gläscher & Adolphs, 2003), and compared the groups on the resulting mean *R* coefficients (after applying Fisher’s *Z* transformation) that reflected the ability to express changes in valence or arousal.

One-way ANOVAs were used to assess group effects on the study dependent variables. Significant group effects were further investigated via two-sided two-sample t-tests (*LSD* corrected). No gender differences were found on the study dependent variables (all *P* ≥ 0.361). Therefore, we report collapsed group data here. We complemented the analysis of AUC by analyses of covariance (ANCOVA) with measurement location (Jena vs. Muenster) as the covariate. Statistical analyses were conducted in IBM SPSS Statistics (v.25, IBM, Armonk, New York, USA). Level of significance was set at *P* = 0.05.

### 2.7. Data availability

The datasets generated during and/or analysed during the present study are partially included in this article (Supplementary Material) or are available from the corresponding author on a reasonable request.

## 3. Results

### 3.1. Baseline variables

Sample characteristics are presented in Table 1. There were no significant differences across groups in terms of age, sex, or education (all *P* ≥ 0.213). Left and right-injured patients did not differ on neuropsychological characteristics (all *P* ≥ 0.289). Visual field examination yielded no evidence for manifest neglect (i.e. no blind spots). On average, total lesion volume (*P* = 0.860) and anatomical distribution of damaged tissue were similar between the patient groups (see Fig.2 for the overlap of reconstructed lesions, Supplementary Fig. 1 for representative anatomical images for each patient, and Supplementary Table 1 for detailed lesion analysis). A significant group difference emerged for depression (*F*[2,34] = 13.30, *P* < 0.001), with both patient groups showing a higher score on the BDI-II compared to the control group (left group vs. control group: *P* < 0.001, right group vs. control group: *P* = 0.003). However, correlative analyses did not reveal any significant association between the BDI-II scores and the study dependent variables (valence and arousal report, AUC). Therefore, BDI-II scores were not included as covariates in the primary study analyses.

**Figure 2.**
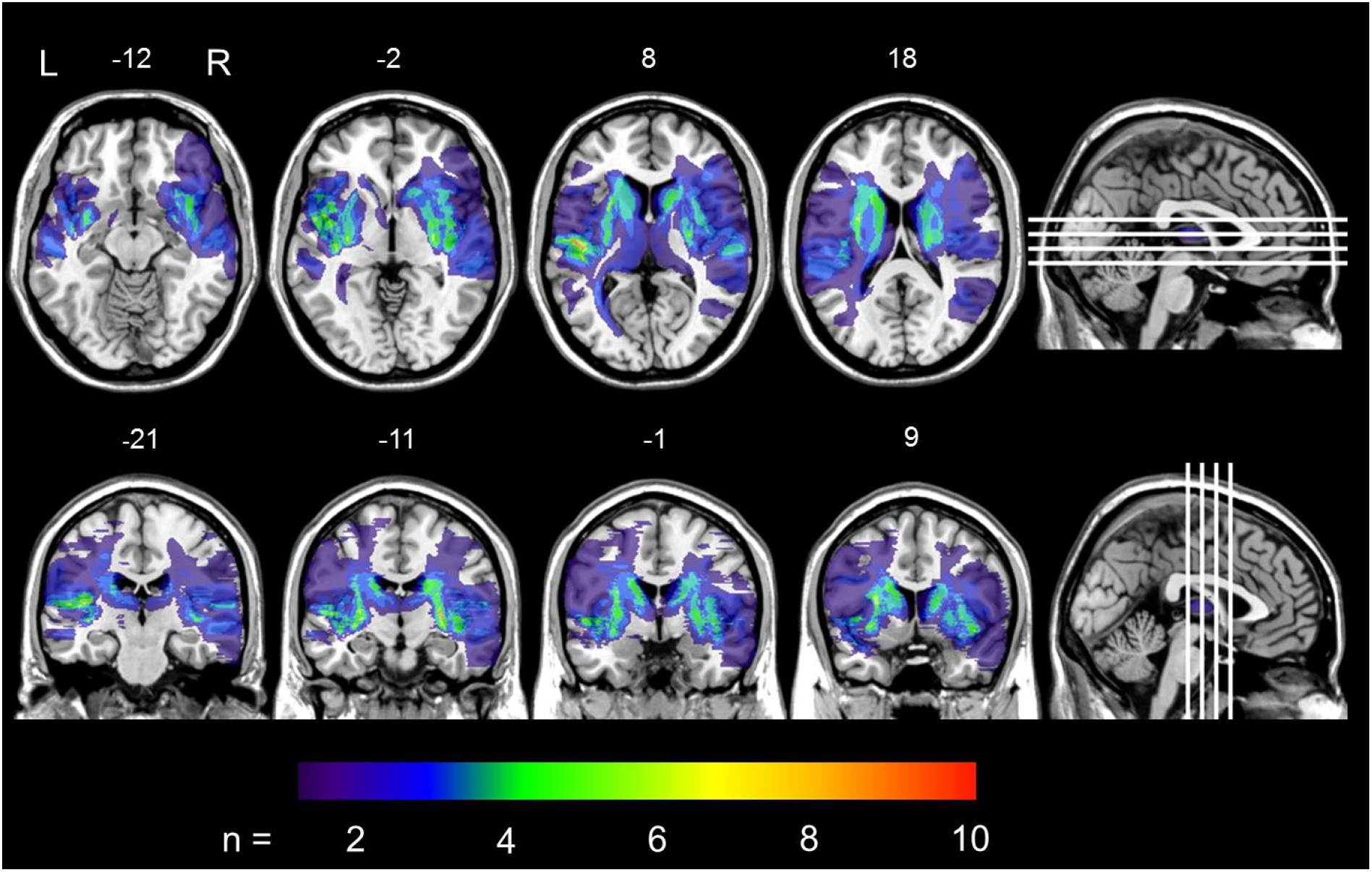
Overlap of lesions within the left (L) and right-lesioned (R) group, overlaid on a T_1_-template in MNI space (1×1×1 mm^3^). MNI coordinates for each axial (z-axis, upper panel) and coronal section (y-axis, lower panel) for visualization are given. The color scale indicates the number of patients with damage in the respective voxel.

### 3.2. Analysis of ratings

We found no effect of Group on mean valence ratings (*F*[2,34] = 0.46, *P* = 0.634) (Fig. 3A), but on mean extremity scores (*F*[2,34] = 7.20, *P* = 0.002) (Fig. 3B). *Post-hoc* comparisons showed significantly lower extremity scores in the right-lesioned group compared to the left-lesioned group (*P* = 0.021) and the healthy control group (*P* = 0.001). A significant effect of Group also occurred for mean arousal ratings (*F*[2,34] = 8.02, *P* = 0.001) (Fig. 3C). *Post-hoc* comparisons revealed significantly reduced arousal ratings in the right-lesioned group relative to the left-lesioned group (*P* = 0.008), and the control group (P < 0.001).

**Figure 3.**
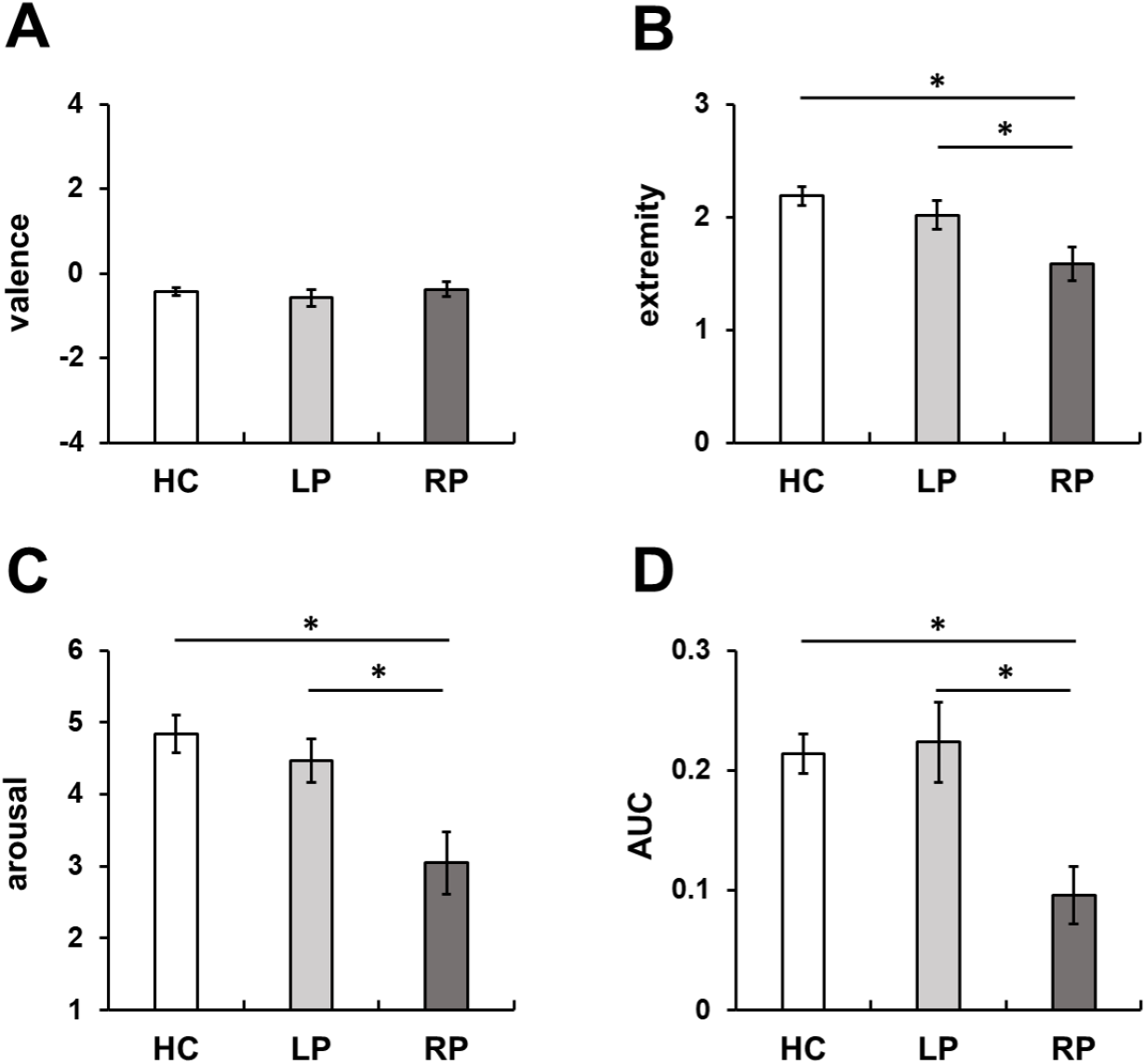
Mean affective ratings (A-C) and autonomic responses (area under the curve, AUC) (D) for each group (HC: healthy control group, LP: left-lesioned group, RP: right-lesioned group). Error bars represent standard error of the mean. * *P* < 0.05, one-way ANOVA for main group effects.

Analysis of *R* coefficients between the subjective and normative valence ratings revealed a significant effect of Group (*F*[2,34] = 5.33, *P* = 0.010) (Fig. 4A). *Post-hoc* tests showed that the right-lesioned group exhibited significantly smaller coefficients than the control group (*P* = 0.003), and marginally smaller coefficients as compared to the left-lesioned group (*P* = 0.094). Analysis of *R* coefficients between the subjective and normative arousal ratings also indicated a significant effect of Group (*F*[2,34] = 10.10, *P* < 0.001) (Fig. 4B). *Post-hoc* analyses showed significantly smaller coefficients in the left (*P* = 0.013) and right-lesioned group (*P* < 0.001) relative to controls.

**Figure 4.**
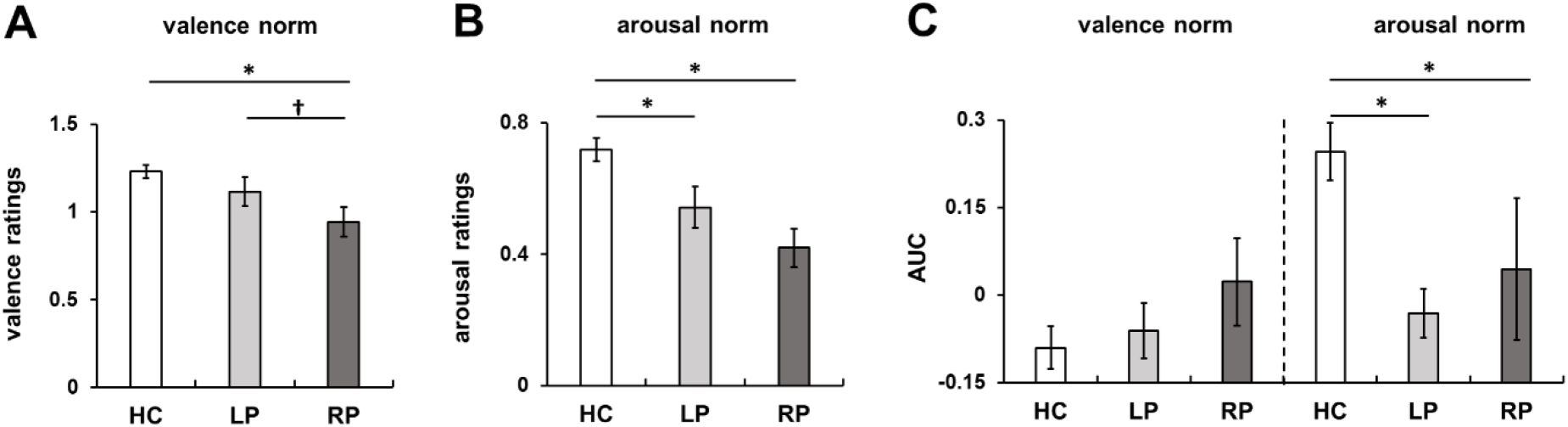
Mean group Fisher Z-transformed correlation coefficients relating normative to subjective valence ratings (A), normative to subjective arousal ratings (B), normative valence and arousal ratings to AUC (area under the curve) (C). HC: healthy control group, LP: left-lesioned group, RP: right-lesioned group. Error bars represent standard error of the mean. * *P* < 0.05, one-way ANOVA for main group effects.

### 3.3. Analysis of autonomic responses

We found a significant effect of Group on mean AUC (*F*[2,31] = 6.38, *P* = 0.005; adjusted for location: *F*[2,30] = 5.61, *P* = 0.009) (Fig. 3D). *Post-hoc* analyses indicated significant reductions of autonomic arousal in the right-lesioned group, as compared to the left-lesioned group (*P* = 0.003; adjusted for location: *P* = 0.029), and the control group (*P* = 0.003; adjusted for location: *P* = 0.001).

Analysis of *R* coefficients between the AUC and normative valence ratings revealed no significant effect of Group (*F*[2,31] = 1.24, *P* = 0.305; adjusted for location: *F*[2,30] = 1.10, *P* = 0.346) (Fig. 4C). However, analysis of *R* coefficients between the AUC and normative arousal ratings indicated a significant effect of Group (*F*[2,31] = 9.36, *P* = 0.001; adjusted for location: *F*[2,30] = 9.93, *P* < 0.001) (Fig. 4C), with significantly smaller coefficients in the left-lesioned (*P* < 0.001; adjusted for location: *P* = 0.001) and right-lesioned group (*P* = 0.011; adjusted for location: *P* = 0.028), relative to the control group.

## 4 Discussion

The present study was carried out to examine how lateralized insular lesions affect autonomic responses and ratings of emotional valence and arousal to standardized emotional scenes. To meet this aim, we systematically compared profiles of SCRs and the cognitive evaluation of stimulus valence and arousal in patients with left or right insular damage and healthy controls. We found a significant reduction in overall SCRs, arousal ratings, and extremity scores in right-lesioned patients, as compared to left-lesioned patients and healthy controls. Additional analyses of correlations between subjective arousal ratings resp. SCRs and normative arousal ratings showed that both lesion groups had evaluative and physiological difficulties to identify changes in stimulus arousal. Although no group differences emerged on overall valence ratings, analysis of correlations between subjective and normative valence ratings displayed reduced accuracy in right-lesioned patients, as compared to left-lesioned patients and healthy controls. Together, our data suggest an idiosyncratic involvement of the left and right insula in emotional processing, with a role of the right insula as integral node for sympathetic arousal and cognitive-affective processing.

The pattern of attenuated autonomic responses and arousal ratings to emotional stimulation in right-lesioned patients points to a direct role for the right insula in the processing of stimulus arousal. The insula is known to subserve interoception and awareness of internal bodily states (Craig, 2014; Critchley, 2005), and has recently been identified as a kind of read-out node that raises awareness of current arousal processes (Damasio, 2018). Our findings resonate with previous reports of right insular involvement in the regulation of the sympathetic tone (Beissner et al., 2013; Craig, 2005; Oppenheimer et al., 1992). Arousal level and the activity of the sympathetic nervous system are closely connected (Critchley, 2005). Thus, by registering deviations from the homeostatic state of sympathetic activity and by perceiving the deviation, the arousal level in the right insula can be recognized (Terasawa et al., 2015). Accordingly, right insular activation has been shown to predict greater magnitude or frequency of the SCR (Critchley, 2009). Patients with right insular lesions often exhibit dysregulation of sympathetic activity to emotional stimulations, or display general hypoarousability (Ibañez et al., 2010; Kim et al., 2017; Manes et al., 1999). In contrast, the left insula seems to support the modulation of the parasympathetic tone (Craig, 2005; Guo et al., 2016; Oppenheimer et al., 1996), and this may explain the preserved physiological and evaluative arousal responses in the left-lesioned group. Importantly, our analyses of correlations between experimental arousal measures and normative arousal ratings show that both patient groups were impaired in their ability to express changes in stimulus arousal in terms of SCRs, or arousal ratings. This finding outlines that both left and right insulae are necessary for adequate cortical representation, control and regulation of autonomic tone, though their roles are different (Craig, 2005). Indeed, integrity of emotional processing relies on the coordinated opponent inhibition between the hemispheres (Craig, 2014).

We found no significant group differences on valence ratings, although our correlational analyses revealed that right-lesioned patients were somewhat less accurate in rating stimulus valence than left-lesioned patients, and controls. However, right-lesioned patients exhibited significantly lower extremity scores than left-lesioned patients, or controls. This finding is in line with a previous clinical report of overall reduced valence ratings for positive as well as negative stimuli after insular lesions (Berntson et al., 2011). There is evidence that interoceptive sensitivity is closely connected with emotional experience (Gu et al., 2013; Lamm & Singer, 2010). Thus, right insular lesions may not disrupt the representation of emotional valence per se, but by disrupting the representation of emotional arousal, rather lead to blunted (ergo less extreme) ratings of emotional valence. Note that emotional blunting is a frequent symptom in patients with right insular dysfunction (Gainotti, 2019).

Previous lesion studies have shown attenuated emotional sensitivity particularly for negative emotions in single patients with right insular dysfunction (Couto et al., 2015; Terasawa et al., 2015; Tippett et al., 2018). Negative emotions are plotted closely on the valence dimension, but they may differ in terms of their arousal level. Right insular lesions may impair interoception that underlies the recognition of arousal level, and may lead to impaired discrimination between negative emotions. Note, however, that disgust seems to rely more on the left insula (Borg et al., 2013; Calder et al., 2000; Holtmann et al., 2018; Kipps et al., 2007; Kumfor et al., 2013; Papagno et al., 2016). Activation of the parasympathetic nervous system is essential in initiating disgust responses (Levenson, 2014). Thus, left-lateralized disgust processing may be due to the left-focused parasympathetic representation.

There are limitations to be noted. The major caveat is the small number of patients, a limitation imposed by the rarity of patients with circumscribed insular lesions. Moreover, we investigated patients with extended lesion areas, which aggravates definite inferences on the neural underpinnings of valence and arousal processing. Lateralization has been mostly documented for the insula. Yet, the described effects may be imparted from insular interconnections with the basal ganglia (Adolfi et al., 2017; Couto et al., 2013; Lewis et al., 2010) rather than the insula itself. The basal ganglia support the recognition of emotion, response/approach motivation, and appropriate emotional behavior (Pierce & Péron, 2020). Furthermore, arousal deficits have also been proposed after damage in other brain areas, including the amygdala (Gläscher & Adolphs, 2003; Holtmann et al., 2018). In particular, the right amygdala contributes to global autonomic activation (Gläscher & Adolphs, 2003). Given the structural and functional complexities of the amygdala and the insula, as well as the rich interconnections between them, further studies are needed to refine distinct neural networks contributing to valence and arousal processing and the specificity of brain regions. Nevertheless, our results suggest hemispheric lateralization as an expedient basis for the investigation of functional involvement of single nodes and their connections within and between hemispheres, for example by applying dynamic causal modelling. Here, effects of insular sub-regions on emotional functions may deserve particular interest (Craig, 2014; Gu et al., 2013; Vicario et al., 2017).

Concluding, this study revealed a disproportionate impairment in arousal processing after right, as compared to left insular lesions. Thus, based on this outcome, we argue for a distinct hemispheric involvement of the insula in emotional processing, potentially based on asymmetrical representations of peripheral autonomic nervous activity in the human forebrain. Our data support the role of the right insula as an integral node for sympathetic arousal and cognitive-affective processes. These results may have relevance for the assessment and treatment of neuropsychiatric disorders that are, at least partly, characterized by insular dysfunction (e.g. Huntington’s disease, or Parkinson’s disease).

## Acknowledgements

All authors appreciate the patients’ contribution to this work and would like to thank them and their families for participation in this study.

## Contributions

TS, WHRM and OH designed the study. CP, WS, NM and CM assessed MRI/ CT scans. OH, MF and II collected the data. OH, IS and JGT analysed the data. OH wrote the first draft of the manuscript.

