## Supplementary Material for "Lateralized deficits in arousal processing after insula lesions: behavioral and autonomic evidence"

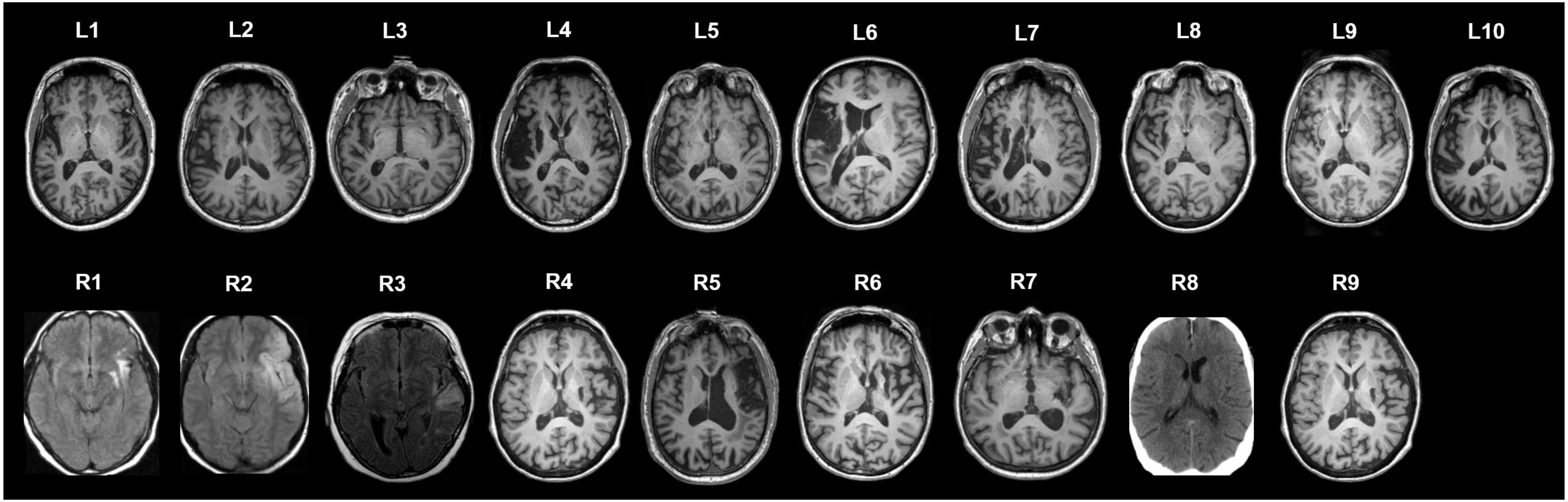


**Supplementary Figure 1:** Overview of representative individual structural images obtained in left-lesioned (L) and right-lesioned (R) patients. $T_1$-weighted axial MR images were obtained in 15 patients, $T_2$FLAIR MR images in three patients (R1, R2 and R3) and CT images in one patient (R8). All images are in neurological orientation (i.e. right on right).

**Supplementary Table 1: Comparative overview of regional lesion volume**

| region | left lesions<br>*n* = 10 | right lesions<br>*n* = 8 | *P* |
|---|---|---|---|
| insula | 2.9 (1.1) | 4.2 (1.7) | 0.534 |
| basal ganglia | 5.1 (1.6) | 6.6 (2.0) | 0.579 |
| white matter tracts adjacent to insula and basal ganglia | 4.0 (1.2) | 3.9 (1.2) | 0.958 |
| thalamus | 1.1 (0.6) | 0.8 (0.4) | 0.644 |
| limbic structures | 0.2 (0.1) | 0.2 (0.1) | 0.831 |
| temporal lobe | 6.4 (1.8) | 10.4 (6.0) | 0.529 |
| central regions | 3.1 (1.9) | 2.5 (1.6) | 0.801 |
| frontal lobe | 4.9 (3.0) | 6.3 (4.3) | 0.785 |
| parietal lobe | 4.7 (3.1) | 1.7 (1.1) | 0.407 |
| occipital lobe | 0.3 (0.3) | 0.01 (0.01) | 0.344 |
| white matter tracts | 3.9 (2.0) | 3.0 (1.5) | 0.729 |
| unclassified tissue | 4.9 (2.4) | 6.2 (3.2) | 0.741 |

*Note:* Data are means (standard error of mean). For each anatomical subdivision, lesion volume is reported in $cm^3$. Standard atlases (AAL atlas, JHU white matter atlas) were used to determine lesion localization and lesion volume on CT and MRI images normalized to MNI space. A claustrum region of interest was manually added to the analyses, since claustrum is not included in the AAL atlas. Basal ganglia damage covered damage to the caudate nucleus, putamen, pallidum and claustrum. Damage of limbic structures comprised damage to the hippocampus, amygdala, cingulate gyrus and parahippocampal gyrus. Temporal lobe damage referred to damage to the temporal pole, superior, transverse, middle and inferior temporal gyrus. Analyses of damage to central regions, frontal, parietal and occipital lobes followed the anatomical parcellation proposed by (Rolls et al., 2015). Damage to external and internal capsule, anterior corona radiate, uncinate and superior fronto-occipital fasciculus was subsumed under damage to white matter tracts adjacent to insula and basal ganglia. Damage to all other deep white matter structures included in the JHU white matter atlas was subsumed under damage to white matter tracts. Unclassified tissue refers to damaged regions that were not included in the atlases.
Statistical comparisons: *t* tests (uncorrected for multiple comparisons).

Rolls, E. T., Joliot, M., & Tzourio-Mazoyer, N. (2015). Implementation of a new parcellation of the orbitofrontal cortex in the automated anatomical labeling atlas. *NeuroImage*, *122*, 1–5. https://doi.org/10.1016/j.neuroimage.2015.07.075
